# Deletion of Glyoxalase 1 exacerbates acetaminophen-induced hepatotoxicity in mice

**DOI:** 10.1101/2023.12.21.572856

**Authors:** Prakashkumar Dobariya, Wei Xie, Swetha Pavani Rao, Jiashu Xie, Davis M. Seelig, Robert Vince, Michael K. Lee, Swati S. More

**Author notes:** Contributed equally (Prakashkumar Dobariya, Wei Xie).

## Abstract

Acetaminophen (APAP) overdose triggers a cascade of intracellular oxidative stress events culminating in acute liver injury. The clinically used antidote, N-acetylcysteine (NAC) has a narrow therapeutic window and early treatment is essential for satisfactory therapeutic outcome. For more versatile therapies that can be effective even at late-presentation, the intricacies of APAP-induced hepatotoxicity must be better understood. Accumulation of advanced glycation end-products (AGEs) and consequent activation of the receptor for AGEs (RAGE) are considered one of the key mechanistic features of APAP toxicity. Glyoxalase-1 (Glo-1) regulates AGE formation by limiting the levels of methylglyoxal (MEG). In this study, we studied the relevance of Glo-1 in APAP mediated activation of RAGE and downstream cell-death cascades. Constitutive Glo-1 knockout mice (GKO) and a cofactor of Glo-1, ψ-GSH, were employed as tools. Our findings show elevated oxidative stress, activation of RAGE and hepatocyte necrosis through steatosis in GKO mice treated with high-dose APAP compared to wild type controls. A unique feature of the hepatic necrosis in GKO mice is the appearance of microvesicular steatosis as a result of centrilobular necrosis, rather than inflammation seen in wild type. The GSH surrogate and general antioxidant, ψ-GSH alleviated APAP toxicity irrespective of Glo-1 status, suggesting that oxidative stress being the primary driver of APAP toxicity. Overall, exacerbation of APAP hepatotoxicity in GKO mice suggests the importance of this enzyme system in antioxidant defense against initial stages of APAP overdose.

## 1. Introduction

The relationship of local redox state with the progression of inflammatory, metabolic and proliferative processes in liver disorders is well-known [1]. Insufficient antioxidant capacity relative to the magnitude of oxidative insults leads to undesirable damage to major biomolecules: protein, lipids and DNA. Cellular antioxidant protection is provided by several enzymes (e.g., superoxide dismutase (SOD), catalase, and glutathione-dependent enzyme pathways) and endogenous molecules (e.g. glutathione (GSH), vitamin E, ascorbic acid). These entities neutralize oxidative insults such as reactive oxygen species (ROS) produced by metabolic processes and the mitochondrial electron transport chain under normal homeostasis. Increased oxidative stress also leads to excess of reactive dicarbonyl metabolites such as methylglyoxal (MEG), a byproduct of glycolysis, also results in ROS generation and oxidative stress. Indeed, hyperglycemia associated with metabolic disorders has been shown to increase MEG levels, resulting in harmful effects on liver function [2]. At higher concentrations, MEG exerts cytotoxic effect by covalently modifying proteins, nucleotides and phospholipids. Alterations in protein structure and function are caused by MEG-dependent formation of “advanced glycation end products (AGE)” and are considered a causative factor behind the resultant “dicarbonyl stress” [3,4]. Important AGEs derived from MEG are the non-fluorescent products 5-hydro-5-methylimidazolone (MG-H1) and tetrahydropyrimidine (THP), as well as the fluorescent argpyrimidine. The AGEs bind and activate the receptor for advanced glycation end products (RAGE). RAGE is expressed in hepatocytes and hepatic stellate cells. The consequence of its interaction with RAGE ligands is activation of various fibrotic, angiogenic, and apoptotic signal transduction pathways and expression of inflammatory mediators such as TNF-α and IL-6, which activate hepatic stellate cells [5–7].

The glyoxalase system is the major enzymatic defense system involved in the regulation of oxidative stress and neutralization of ROS resulting from dicarbonyl MEG. This system consists of two crucial enzymes glyoxalase-1 (Glo-1) and glyoxalase-2 (Glo-2). Glo-1 is a dimeric metalloprotein containing a Zn^+2^ ion in each of the two subunits [4]. Human Glo-1 expression is controlled by the activator protein-2α (AP-2α), nuclear transcription factor-κB (NF-κB), early gene 2 factor isoform 4 (E2F4), antioxidant-response element (ARE), nuclear factor erythroid-2-related factor 2 (Nrf2), among other entities [4]. Glo-1 is an aldoketomutase—it rearranges the spontaneously formed hemithioacetal of GSH with MEG into S-D-lactoylglutathione. The latter is hydrolyzed by Glo-2 into relatively non-toxic D-lactate with liberation of GSH [8]. Glo-1 also reduces MEG formed by other metabolic processes like, threonine catabolism, ketone body metabolism and degradation process of glycated protein [3,8]. Decreased levels of Glo-1 and increased intracellular dicarbonyl stress occurs in liver fibrosis, cirrhosis, non-alcoholic steatohepatitis (NASH) and hepatocellular carcinoma. Pharmacological modulation of Glo-1 as well as silencing of RAGE have shown promise in preclinical models of CCl_4_-induced liver fibrosis and hepatocellular carcinoma, respectively [9]. Glo-1 mediated detoxification of MEG and structurally related dicarbonyls is considered a rate-limiting step [3]. Therefore, modulation of the Glo-1/AGE/RAGE axis for management of chronic liver diseases is explored by us and many others in the field.

Although the role of Glo-I and (R)AGE in chronic liver disease has been described [3,9], its role in acute liver failure is relatively understudied. Autoimmune disease, hepatitis, toxins and drugs could cause acute liver failure. Drug-induced liver injury contributes to nearly 60% of those cases. Among these, acetaminophen (N-acetyl-p-aminophenol, APAP)-induced liver toxicity is the leading cause of drug-induced acute liver failure in the United States [10]. An estimated 50,000 people in the United States are hospitalized annually due to APAP overdose, and up to 500 people die each year. APAP overdose results in accumulation of high levels of the extremely reactive metabolite, N-acetyl-p-benzoquinoneimine (NAPQI) that rapidly depletes cellular GSH levels, induces oxidative stress and accumulation of AGEs, ultimately causing cell damage and death. N-acetylcysteine (NAC), a GSH precursor, is a classical antidote for the APAP-induced hepatotoxicity used in the clinic. Mechanistically, NAC triggers GSH synthesis, enhances glutathione-S-transferase (GST) activity, and facilitates detoxification by scavenging ROS. Reduced levels of GSH after APAP overdose are reported in models of APAP toxicity and also in patient serum [11,12]. Being the cofactor for Glo-1 enzyme, deficiency of GSH caused by high-dose APAP is expected to perturb the Glo1/AGE/RAGE axis, thus increasing oxidative stress and promoting hepatocyte necrosis. It is therefore essential to study the effects of GSH depletion and Glo-1 inactivation on APAP-induced liver toxicity. Pharmacological modulation of Glo-1 as well as gene deletion are expected to shed light on the complex interplay between these pathways involved in hepatotoxicity and provide insights into potential therapeutic interventions for APAP overdose.

Toward that goal, we utilized constitutive Glo-1 knockout mice to decipher the role of Glo-1 detoxification pathway in APAP-induced hepatotoxicity. In a previous study, we described a bioavailable GSH surrogate, ψ-GSH, as a potential antidote for APAP liver toxicity [13]. Utility of GSH by itself or GSH-precursors as antidotes is limited due to rapid cleavage of GSH by a ubiquitous γ-glutamyltranspeptidase, resulting in poor bioavailability. Designed to address these limitations, ψ-GSH mitigated APAP toxicity and improved survival, even when administered post-APAP overdose —mimicking the clinical scenario in a mouse model of APAP hepatotoxicity. It also serves as a replacement for GSH itself in major GSH-dependent enzymatic pathways [14]. We have employed ψ-GSH to study the central function of the Glo-1 pathway in neurodegenerative pathology and evaluated its potential as a therapeutic for management of neurodegenerative disorders such as Alzheimer’s disease [15,16]. Here, we use ψ-GSH as a pharmacological tool for comprehensive characterization of the Glo-1 pathway in APAP-induced liver toxicity.

In this study, we created an APAP hepatotoxicity model in whole-body Glo-1 knockout mice (Glo-1^−/−^, GKO) and examined the effects of ψ-GSH on this model. Whole-body Glo-1 knockout mice were previously generated and characterized by Jang et al. [17]. Increased liver AGEs and reduced anxiety-like behavior were noted in GKO mice compared to age-matched wild-type mice. We observed exacerbation of liver toxicity induced by high-dose APAP in GKO mice. Histological analysis suggested an unexpected and distinct pattern of hepatocyte necrosis when compared to that in wild-type (WT) mouse: *through steatosis rather than inflammation*. APAP metabolism was also significantly altered in the knockout mice. Further, biochemical analysis confirmed involvement of the Glo-1/AGE/RAGE axis in GKO mice. Intervention with ψ-GSH, due to innate antioxidant potential, was successful in mitigating oxidative stress in both WT and GKO mice. Thus, this study for the first time demonstrates the significance of the Glo-1 pathway in oxidative stress induced by APAP overdose.

## 2. Materials and Methods

### Materials

Acetaminophen (Spectrum Chemical Corp, New Brunswick, NJ, #AC100), triethanolamine (Sigma-Aldrich Corp, # t58300), 2-vinylpyridine (Alfa Aesar, Ward Hill, MA, #A14056), RIPA Buffer (Cayman Chemical Co., Ann Arbor, MI, # 10010263), GSH MES Buffer (Cayman Chemical Co., # 703010), cOmplete Protease Inhibitor Cocktail (Roche, Penzberg, Germany, #4693124001).

### Animals

Male and female C57BL/6 age 18–20 weeks were obtained from Charles River Laboratories and used in the present study. Sperm carrying Glo1^tm1a(KOMP)Mbp^ allele was obtained from the European Mouse Mutant Achieve (EM:09893) and live mice was generated via in vitro fertilization at University of Minnesota Mouse Genetics Laboratory. Resulting heterozygous mice were mated to generate homozygous GloKO mice (GKO) and maintained in the Lee lab at the University of Minnesota. All experimental procedures and animal handling were conducted in accordance with the national ethics guidelines and complied with the Institutional Animal Care and Use Committee (IACUC) protocol requirements of the University of Minnesota, Minneapolis, MN. Every effort was made to minimize animal suffering and the number of animals used in this study. The animals were housed in university facility in groups of four per cage under controlled environmental conditions on a 12 h-12 h light-dark cycle and were allowed access to food and water ad libitum. The experimental procedures were carried out during the light phase of the light/dark cycle.

### Behavioral assessment of GKO mice

Twelve-week-old male and female WT and GKO mice were used for behavior analysis. For open field test (40 cm × 40 cm × 40 cm), mice were habituated for an hour in a dark recording room, after that, the mouse was put in the center of the open field and video recorded for 10 min, and the first 5 min were analyzed. For light-dark box test, after habituating for an hour in the test room, each mouse was placed in the center of the lighted zone, facing toward the maze wall, which was recorded for 15 min, and the first 10 min were analyzed. For tail-flick test, mice were habituated for an hour in recording room. Mice tails were exposed to a light beam centered on ventral surface, about 15 mm from the tip of the tail and the latency for withdrawal of the tail is recorded. In T-maze test, spontaneous alternation protocol was used for assessment of working cognitive behavior. After habituation for an hour in the testing room, mice were allowed to freely explore the entire maze. This was followed by confinement in the start arm for 30 s and 15 free-choice trials to calculate percent alternations.

### Animal Handling

The mice were allowed to acclimate to the facility for at least one week prior to the start of the experiment. Prior to drug administration, C57BL/6 and GKO mice were fasted overnight, weighed, and randomly assigned to different treatment groups (N=10-12). Acetaminophen was dissolved in warm saline and injected intraperitoneally (i.p.) at a dose of 250 mg/kg (1.65 mmol/kg). At the designated time, either ψ-GSH (800 mg/kg) or saline was administered to the mice by i.p. route with control groups receiving saline only. Food was returned to the cages after the final drug injection, and animals were monitored every 30 minutes for the first 2 hours. At the end of the experiments, blood and liver tissue samples were collected from the mice at 24 hours after APAP administration. Liver tissues were immediately frozen and stored at −80 °C until further use or fixed in 10% neutral buffered formalin for histological analysis.

### ALT level assessment in serum

Serum alanine aminotransferase (ALT), creatinine and creatinine kinase (CK) measurement was used to assess liver and kidney damage caused by APAP overdose. Blood samples were collected from the mice at 24 hours post APAP overdose. The clotted blood was centrifuged to obtain serum, which was then stored on ice until it was submitted to the Veterinary Diagnostic Laboratory at the University of Minnesota for blood chemistry measurement.

### Histology

Liver morphology was evaluated by histological analysis of H&E-stained liver sections. The liver harvested from the mice at 24 hours post APAP overdose was immediately fixed with 10% formalin. The fixed liver samples were then submitted to the Comparative Pathology Shared Resource at the University of Minnesota for processing.

### APAP and Metabolites in Serum Samples

Mouse serum samples collected as described in ALT assessment were used for measurement of APAP and metabolites by LC-MS/MS analysis. Serum samples were processed by addition of 90 μL of acetonitrile to 10 μL of the serum sample. The samples were mixed by vortexing for 30 seconds and then centrifuged at 21,130xg for 5 minutes at 4 °C. The supernatants were collected and transferred to a fresh 1.5 mL microcentrifuge tube.

Liquid chromatography-tandem mass spectrometry (LC-MS/MS) analysis was conducted using an integrated system consisting of an Agilent 1260 High-Performance Liquid Chromatography (HPLC) device (Agilent Technologies, Santa Clara, CA, USA) coupled with an AB Sciex QTRAP 5500 mass spectrometer (AB Sciex LLC, Toronto, Canada) with slight modifications to the method described previously [18].

Briefly, chromatographic separation of the samples was achieved using a Thermo Aquasil C18 column (150 × 2.1 mm, 3 μm). A binary mobile phase system at a constant flow rate of 0.3 mL/min was employed with mobile phase A consisting of 0.1% formic acid in water and mobile phase B consisting of 0.1% formic acid in acetonitrile. The gradient elution profile began with a linear increase from 0% to 5% B over 2 minutes. Subsequently, the gradient transitioned from 5% to 80% B over another 2 minutes, held at 80% B for 1.8 minutes, and then returned to 0% B over 0.2 minutes. Finally, the column was equilibrated at 0% B for 6 minutes. Only desired fractions of eluates, corresponding to the 1.3 to 10-minute retention time window, were collected for subsequent LC-MS/MS analysis. Samples were introduced into the mass spectrometer via electrospray ionization in a fast polarity-switching mode. The instrument parameters were set as follows: curtain gas at 25 psi, collision-activated dissociation (CAD) gas at medium flow, ion spray voltage at 5000 V or −4500V, source temperature at 650 °C, gas 1 at 60 psi, and gas 2 at 50 psi. For targeted quantitation, MRM was employed by monitoring specific mass transitions for each analyte as confirmed by commercial standard of each metabolite and transitions are listed as follows: m/z 152.1 → 110.1 (APAP); m/z 182.1 → 108.0 (APAP−OMe); m/z 271.1 → 140.0 (APAP−Cys); m/z 313.1 → 208.0 (APAP−NAC); m/z 457.1 → 328.1 (APAP−GSH) in positive mode and m/z 230.0 → 150.0 (APAP−Sulf); m/z 326.0 → 150.0 (APAP−Gluc) in negative mode.

### Measurement of liver GSH content

The GSH:GSSG ratio was determined by following the protocol included in the glutathione assay kit (Cayman Chemical Co., # 703002). Briefly, liver tissues were harvested, rinsed with PBS, and homogenized using 2 mL of 50 mM MES buffer having 1 mM EDTA, pH 6.0 per gram. Supernatant was collected after centrifugation at 10000×g for 15 minutes at 4 °C. Deproteinization was carried out by adding equal amount of 10% metaphosphoric acid. Deproteinized homogenate (100 μL) was neutralized with 5 μL of 4 M triethanolamine solution and diluted 150 times with MES buffer. To measure the reduced GSH (GSSG), deproteinated samples were treated with 1M 2-vinylpyridine solution (10 μL) and allowed to incubate for an hour at room temperature. GSH and GSSG level was determined from the absorbance after reaction with 5-thio-2-nitrobenzoic acid at 412 nm as per the manufacturer’s instructions.

### Lipid Peroxidation assay

Hepatic lipid peroxidation was measured by TBARS assay. The levels of malondialdehyde (MDA), a breakdown product of lipids, were determined colorimetrically using TBARS assay kit (Cayman Chemical Co., Cat. No. 10009055) as described previously.

### Evaluation of the protein carbonyl content

Level of the protein carbonyl present in the liver samples was measured using protein carbonyl colorimetric assay kit (Cayman Chemical Co., Cat. No. 10005020) and Dot Blot assay kit (Abcam, Cat. No. ab178020). For Dot blot analysis, liver homogenates were first derivatized with DNPH per manufacturer’s instructions. Total 50 ng of the derivatized protein was used to determine the level of the protein carbonyl in each group.

### Determination of liver Superoxide Dismutase (SOD) activity

The SOD activity of liver homogenate (containing 50 mM MES, pH 6.5, 1 mM EDTA) was determined using commercial SOD assay kit (Cayman Chemical Co., # 66970752). Briefly, 10 µl of the liver homogenate was treated with 200 µL of radical detector solution provided in the kit. Reaction was initiated by adding 20 µL of xanthine oxidase and the rate of change in the absorbance at 450 nm was used to calculate SOD activity per sample.

### Determination of liver catalase activity

To determine the catalase activity, the procedure described in the catalase assay kit (Cayman Chemical Co., # 66970752) was followed. Briefly, the liver homogenate was treated with 30 µL methanol and the reaction was initiated by addition of H_2_O_2_. The formaldehyde produced was measured by reaction with 4-amino-3-hydrazino-5-mercapto-1,2,4-triazole (Purpald). The complex upon oxidation with potassium periodate produces purple color, which was measured by absorbance at 540 nm to calculate catalase activity of the sample as described in the kit.

### Western blot analysis

Part of livers were lysed with RIPA buffer containing protease inhibitors (Roche). Protein concentrations were determined using the BCA Protein Assay Kit (Thermo Fisher Scientific). The proteins were denatured and equal amounts of protein (a total of 40 µg/lane) from different groups as indicated in the figures were separated by SDS/PAGE and transferred to polyvinylidene fluoride (PVDF) membrane via a wet transfer method for immunoblot analyses. PVDF membrane was blocked for 2 h using 5% fat milk and incubated with antibody to Cleaved caspase-3 (#9661), RIPK3 (#95702), p62 (#5114), RAGE (ab3611), AGE (ab9890), Glo1 (ab137098), Glo2 (AF5944), HMGB1 (3935S), MG-H1 (STA-011), Bax (sc-20067) and antibody to α-tubulin (ab4074) at 4 °C overnight. Primary antibodies were labeled with horseradish peroxidase-conjugated secondary antibody (1:5000) and ECL substrates (Bio-Rad). The densitometric intensities of the bands were quantified by using ImageJ software.

### Statistical Analysis

Statistical analyses were performed using GraphPad Prism ver 10. Data are expressed as the mean ± standard error of the mean. All data are analyzed using one or two-way analysis of variance (ANOVA) with Tukey’s or Sidak’s post-hoc test or student t test, wherever appropriate. Statistical significance was held at p < 0.05.

## 3. Results

### 3.1. Liver toxicity induced by APAP overdose is potentiated in Glo-1 knockout mice

A single high dose of APAP has been used to generate a mouse model that enabled the study of toxicological mechanisms as well as protective interventions. We have previously studied effects of potential antidotes for APAP overdose in fasted Swiss Webster mice [13,18,19]. This study evaluates the role of the Glo-1 pathway in APAP-induced hepatotoxicity, for which we employed constitutive knockout of Glo-1 (Glo-1^−/−^, GKO) in C57BL/6N background. Due to higher susceptibility of C57BL/6N over C57BL/6J mice to APAP-toxicity [20], GKO mice were maintained in 6N background. Behavioral characterization of GKO mice has been reported previously [17]. We confirmed protein expression of Glo-1 and Glo-2 in GKO mice by western blot (Supporting Information, Figure S1) and behavioral phenotype in open field, light-dark box, T-maze, and tail flick tests. Significant behavioral impact of Glo-1 was noted only in male GKO mice in the open field and light-dark box tests, which is in agreement with the previous reports claiming correlation between Glo-1/MEG concentration and anxiety-like behavior [21,22]. Increase in time spent in the center zone of open field box (Supporting Information, Figure S2-A) and in the lit area of the light-dark box (Supporting Information, Figure S2-D), indicators of anxiolytic behavior, were observed in GKO mice. Mice did not show signs of motor deficits based on the total distance moved (Supporting Information, Figure S2-B). No change in the exploratory behavior was seen in the T-maze spontaneous alternation test (Supporting Information, Figure S2-C). Additionally, response to heat stimuli in the tail-flick test did not differ between WT and GKO mice (Supporting Information, Figure S2-D).

We first calibrated a dose of APAP that causes reproducible and measurable liver enzyme elevation, such as serum alanine aminotransferase (ALT), in GKO mice as a gauge of liver toxicity. The dose equivalent to our previous studies (370 mg/kg) created a much higher response, with effect on animal viability. Thus, we selected a 250 mg/kg dose in this study which produced significant ALT elevation without affecting survival. This dose caused more than 120-fold elevation in the ALT level when compared to saline treated animals (WT-APAP, 4906.52 ± 662.16 *vs* WT-saline, 40.00 ± 6.86 U/L; Figure 1A). Dramatic increase in ALT levels was observed in APAP-treated GKO mice (7919.80 ± 799.07 U/L) over the matched saline-treated controls (56.33 ± 16.48 U/L) and most noticeably, over the WT-APAP treated mice (1.6-fold). Additionally, change in the ALT level was significant only in male mice compared to female mice. Treatment with ψ-GSH showed significant hepatoprotection by alleviating the increased ALT levels induced by APAP overdose in WT, as previously reported by us in an independent study [13] and GKO animals (Figure 1B). Designed as a metabolically stable analog of GSH, the inherent antioxidant nature of ψ-GSH due to aliphatic thiol is capable of non-enzymatic neutralization of ROS generated by APAP and could explain hepatoprotection offered by ψ-GSH in APAP-treated GKO and WT mice.

**Figure 1.**
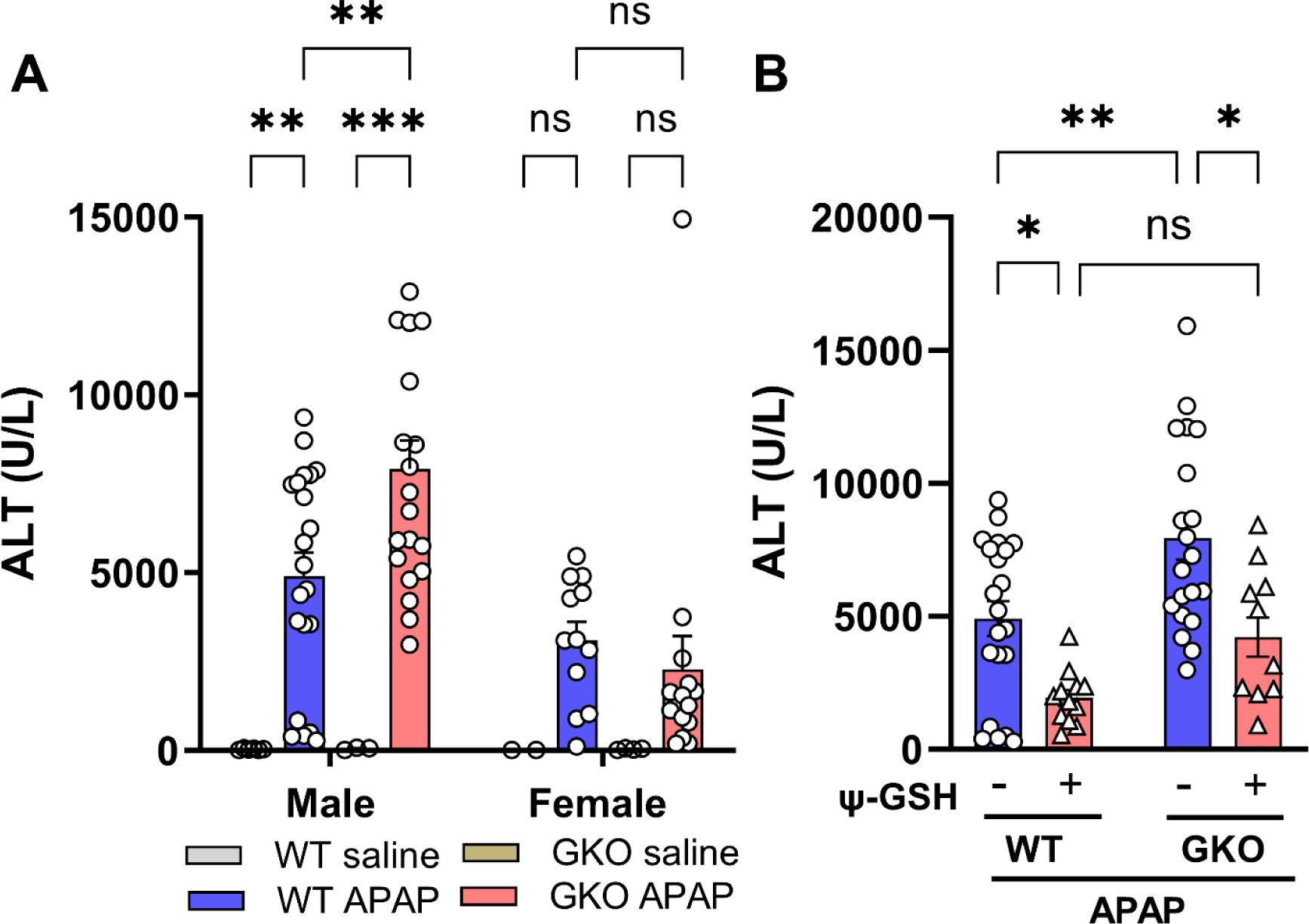
The effect of Glo-1 deletion on hepatotoxicity induced by high dose APAP. (A) Serum samples were collected 24 hours after intraperitoneal administration of APAP (250 mg/kg) from C57BL/6 mice to determine ALT levels. Significant elevation of serum ALT levels was noticed in male GKO mice compared to WT mice, while female mice were resistant to this APAP-induced ALT elevation. (B) Serum ALT levels in mice pretreated with ψ-GSH (800 mg/kg, i.p.) 30 min prior to injection of APAP (250 mg/kg, i.p.). Significant protective effect of ψ-GSH treatment was observed in WT-APAP and GKO-APAP groups (* p < 0.05; ** p < 0.01; *** p < 0.001; ns not significant; two-way ANOVA followed by Tukey’s post-hoc multiple comparison test).

Histopathological evaluation of liver tissue (Figure 2) corroborated the gender-specific findings in ALT elevation. Female mice treated with high-dose APAP showed low-to-moderate necrosis (score 1-2) of the hepatocytes irrespective of the Glo-1 status. In male mice, the WT-APAP group showed coagulative necrosis (score 3-4) covering 30-60% of the centrilobular to midzonal area with markedly swollen hepatocytes (inflammation score 2-3), as reported by others [23,24]. APAP treated GKO mice, in addition to necrosis (score 3-4) and swollen hepatocytes (inflammation score 1), exclusively showed intracytoplasmic, empty to pale pink discrete vacuoles, indicative of microvesicular lipid accumulation (Figure 2, Inset, bottom row). Such microvesicular steatosis, a type of hepatocellular degeneration, was severe in the centrilobular zone of APAP treated GKO mice (score 3-4). Previously, Spanos et al. [25] have also reported the dramatically increased level of intracellular lipids and MEG accompanied by decreased Glo-1 protein expression in HepG2 cells treated with oleic acid [25]. Additionally, reduced reservoir of the GSH due to lack of Glo-1 expression makes steatotic hepatocytes more susceptible towards APAP-induced hepatotoxicity [26]. Treatment with ψ-GSH effectively reduced the extent of necrosis and inflammation in both WT and GKO mice.

**Figure 2.**
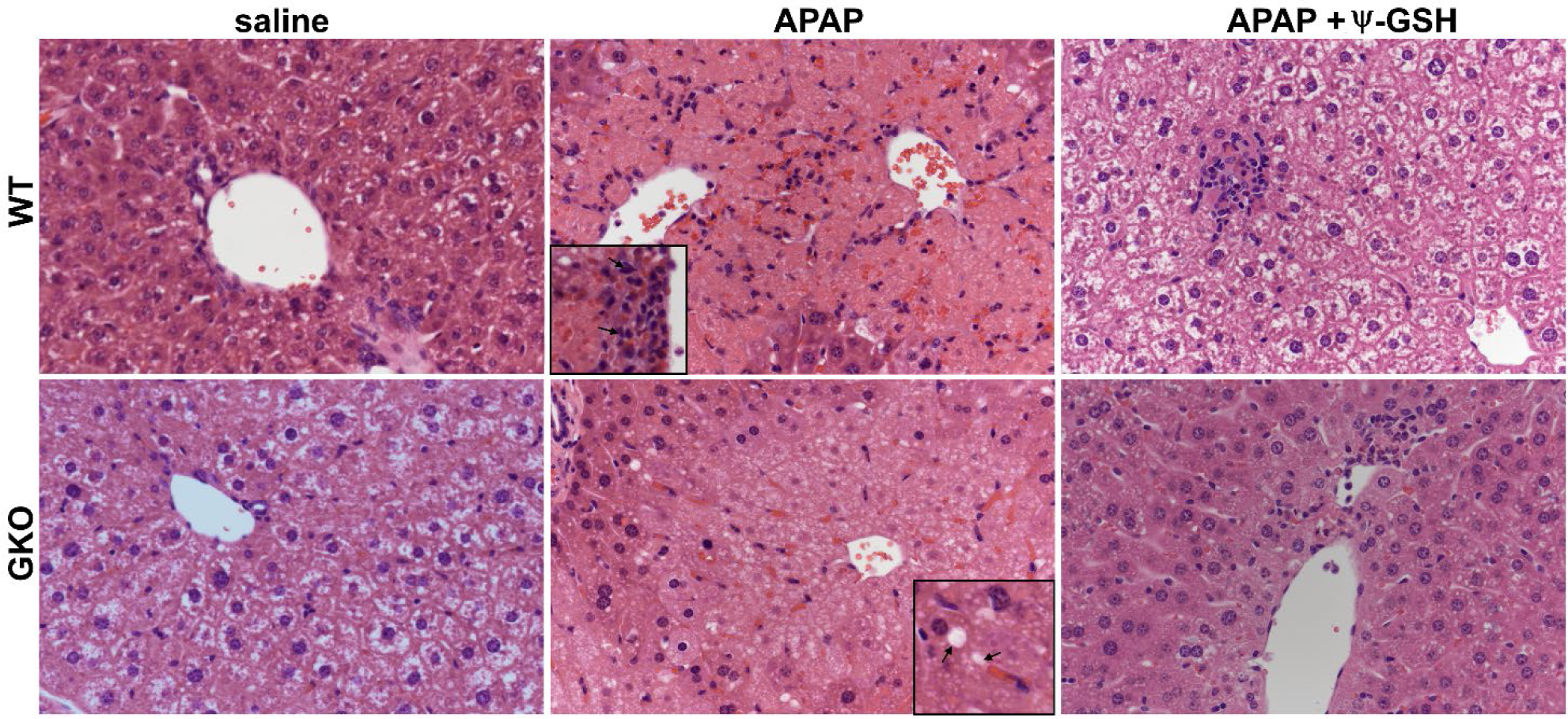
Histological examination of the liver tissue from WT and GKO mice exposed to APAP overdose. Saline treated WT and GKO mouse livers showed intracytoplasmic vacuoles (clear cytoplasm, physiological). APAP treatment resulted in coagulative necrosis in centrilobular zone with similar extent of necrosis in both WT and GKO mice. Inflammatory cells were observed surrounding the central vein in WT-APAP mice (inset, top row), while necrotic hepatocytes showed prominent microvesicular lipid accumulation in GKO-APAP group (inset, bottom row). Ψ-GSH treatment reversed histological changes caused by APAP overdose.

To investigate lipid changes in GKO mice treated with APAP, we examined serum cholesterol and triglyceride levels in these treatment groups. No significant differences were noted in WT and GKO mice in the presence and absence of APAP (Supporting Information, Figure S3), However, noticeable increase in creatinine (∼5.7-fold) and creatinine kinase (∼2.3-fold) was observed in GKO-APAP mice over APAP treated WT mice. This observation is in line with prevalence of chronic kidney disease in patients with hepatic steatosis, which shows association of serum creatinine with hepatic steatosis risk [27]. Exacerbation of intracellular lipid accumulation are proposed to contribute independently toward non-alcoholic fatty liver disease and chronic kidney disease [28]. These observations collectively indicate exacerbation of APAP hepatotoxicity in GKO mice compared to WT animals in a degenerative manner by a unique mechanism involving hepatocellular steatosis.

### 3.2. Deletion of Glo-1 reduced overall metabolic detoxification of APAP

At the therapeutic dose, majority of administered APAP is metabolized by phase-II metabolism by conjugation to glucuronide and sulfate metabolites [29]. A small portion (5-10%) is oxidized by microsomal CYP 2E1 and CYP 1A2 to a highly reactive intermediate, N-acetyl-p-benzoquinone imine (NAPQI) [30]. Hepatic GSH effectively quenches the toxic NAPQI under normal homeostasis. However, under disease conditions or accidental overdose wherein the supply of GSH is limited, NAPQI exerts toxic effects by irreversibly modifying cysteine residues in cellular proteins, exacerbating the oxidative stress and resulting cellular damage [13,31–34]. To understand the effect of Glo-1 on APAP toxicity, we examined the effect of Glo-1 deletion on the levels of major metabolites of APAP. Concentrations of individual APAP metabolite in serum samples were determined using LC-MS/MS (Figure 3 and Supporting Information, Table 1). Serum APAP levels were similar in male WT and GKO mice (Figure 3A). A trend toward higher intact APAP levels was noted in female WT mice, while it was significant in GKO-APAP mice compared to GKO-saline group. APAP-Glucuronide concentrations were similar in male WT and GKO mice; however, were elevated in respective female mice with highest levels found in GKO-APAP group (Figure 3B). The levels of APAP-sulfate were significantly (Figure 3C) reduced in GKO mice compared to WT mice. However, the effect was significant in male mice, while the female mice showed higher APAP-sulfate levels, partly explaining resistance of female mice toward APAP toxicity. Possibly due to higher oxidative stress and reduced GSH levels, conjugation of APAP to GSH, or cysteine protein thiols (phase II) was significantly reduced in the GKO male mice. Thus, resultant levels of non-toxic APAP metabolites such as APAP-GSH, APAP-Cys, and APAP-NAC conjugates were markedly decreased over those in APAP-WT mice (Figure 3D, 3E, and 3F). Furthermore, reduced APAP hydroxylation (phase I) followed by phase II APAP methylation was observed in GKO mice, although the absolute levels of APAP-OMe were significantly lower in both WT and GKO mice when compared to other APAP metabolites (data not shown). These results indicate ineffective detoxification of APAP in the absence of functional Glo-1, contributing toward toxicity of the parent APAP. Table 1 lists concentrations of individual APAP metabolites in WT and GKO mice.

**Figure 3.**
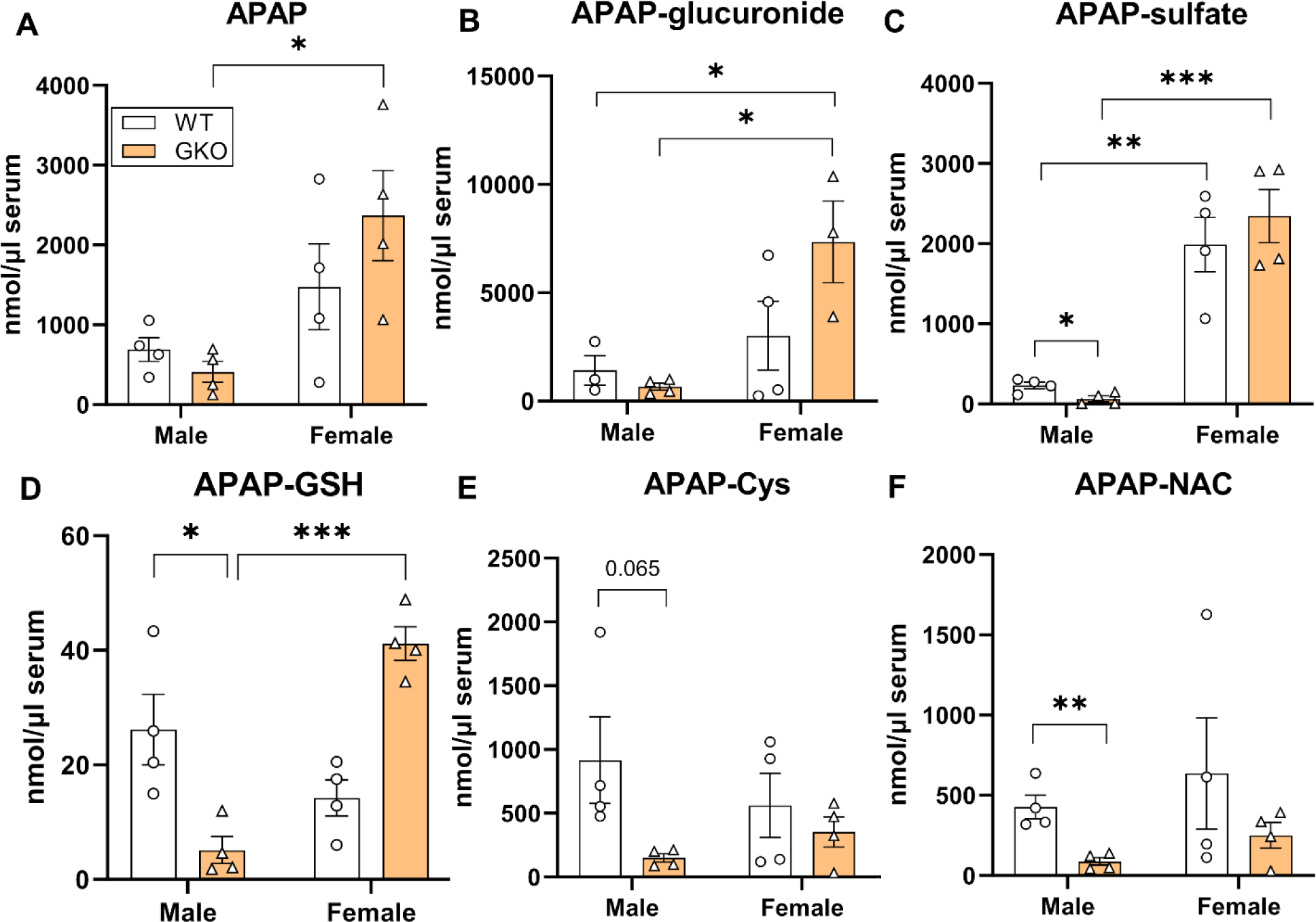
Effect of Glo-1 knockout on the levels of APAP and its metabolites in serum. Serum samples from fasted C57BL/6 mice collected 24 hours after administration of APAP overdose were subjected to metabolite identification by LC-MS/MS. Concentrations of each APAP metabolite from WT and GKO mice are displayed as mean ± SEM. (A) APAP, (B) APAP cysteine adduct (APAP-Cys), (C) APAP N-acetylcysteine adduct (APAP-NAC), (D) APAP glutathione adduct (APAP-GSH), (E) APAP sulfate (APAP-sulfate), (F) 3-methoxy APAP (APAP-OMe). (* p<0.05; **, p<0.01; ***, p<0.001; two-way ANOVA followed by Tukey’s post-hoc test).

### 3.3. Increased oxidative stress observed in APAP-treated Glo-1 KO mice

Oxidative stress has been widely reported as a factor responsible for liver necrosis upon APAP overdose in multiple animal models. As glyoxalase is the major pathway for detoxification of oxidized sugar metabolites and is intricately involved in the oxidative stress mediated by AGE formation. It is thus expected that inherent stress produced by deletion of Glo-1 would be compounded by high-dose APAP. Alteration in biochemical parameters of oxidative stress such as GSH levels, protein carbonyl content, lipid peroxidation and activities of antioxidant enzymes like superoxide dismutase (SOD), catalase, were used as indices of liver oxidative stress. Due to clear hepatotoxic phenotype offered by male mice, only male mice were analysed for changes in biochemical parameters. Similar to our previous reports, we observed attenuation of redox potential in the livers of APAP-treated WT mice (Figure 4A). Surprisingly, GKO-saline mice showed significantly higher redox (GSH:GSSG) ratio compared to WT-saline (GKO-saline, 7.20 ± 0.86 *vs* WT-saline, 4.29 ± 0.64). This could be attributed to a compensatory mechanism exhibited by GKO mice to overcome the non-functional Glo-1 pathway [35,36]. APAP treatment reduced GSH/GSSG ratio in WT and GKO mice, which was reversed significantly in WT group by ψ-GSH (p < 0.01, *t*-test). Biochemical analysis of lipid peroxidation (Figure 4B) showed significantly higher malonaldehyde (MDA), one of the final products of fatty acid oxidation, in APAP treated mice. This increase was higher in APAP-treated GKO mice compared to WT mice (GKO-APAP, 0.57 ± 0.05 *vs* 0.41 ± 0.04 μM/mg protein; Figure 3B), which corroborated with microvesicular steatosis observed in these mice indicative of imbalance in lipid biology. Similar trend was also observed in protein carbonyl content (Figure 4C, 4E) in APAP treated animals by biochemical assay (DNPH assay) as well as by dot blot analysis. A trend toward increased protein carbonyls in GKO-APAP group compared to WT-APAP animals was apparent (p < 0.05, *t-*test). Given the compensatory GSH mechanism exhibited by GKO mice, we decided to study the effect of APAP-overdose on non-GSH dependent antioxidant enzymes. Changes in the activity of important oxidoreductase enzymes like SOD and catalase, which play crucial role in antioxidant defence against APAP-induced oxidative stress, were examined. Compared to saline-treated mice, APAP-treated WT and GKO mice showed lower SOD activity (∼1.6 fold over respective saline groups, Figure 4D), while almost similar catalase activity (Supporting Information, Figure S4) in all treatment groups. Intraperitoneal administration of ψ-GSH effectively mitigated the emergent oxidative stress in APAP-treated WT mice. This was evident from elevation of the redox (GSH:GSSG) ratio, reduced lipid and protein oxidation levels, and increased SOD activity. These results reproduced our previous observations from Swiss Webster mice [13], demonstrating the ability of ψ-GSH in easing the oxidative stress induced by high-dose APAP. The protection offered by ψ-GSH in GKO mice using these indices of oxidative stress was almost equal to WT mice, suggesting major contribution of Glo-1 independent antioxidant mechanisms.

**Figure 4.**
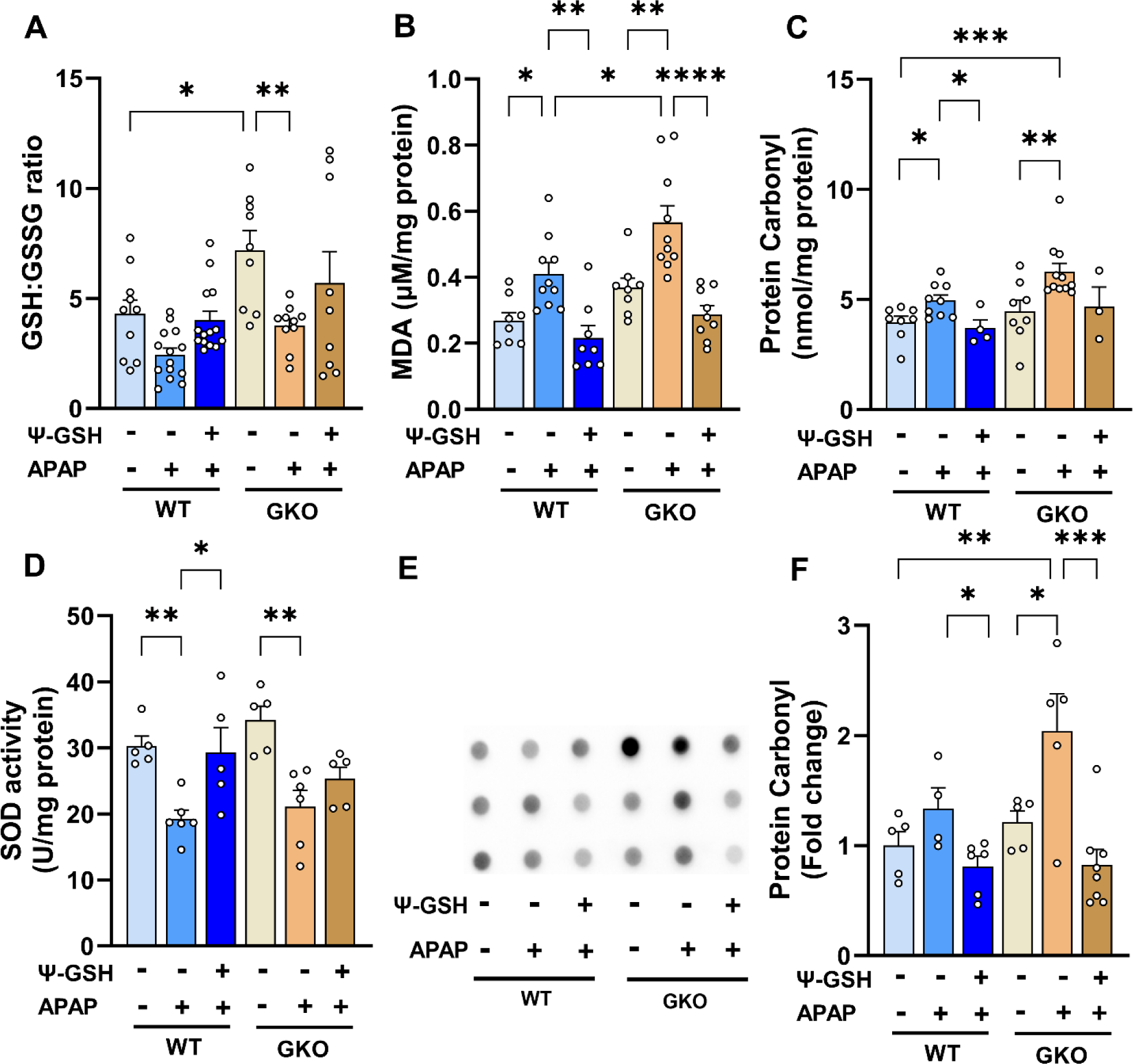
The effect of Glo-1 deletion on the levels of liver oxidative stress markers in APAP-overdosed mice. All liver tissues for oxidative stress assay were collected at 24 h post overdose of APAP (250 mg/kg, i.p.). ψ-GSH (800 mg/kg, i.p.) was administered 30 min prior to APAP injection. (A) liver GSH:GSSG ratio, (B) liver malondialdehyde levels (MDA μM per mg protein), (C) liver protein carbonyl (nmol per mg protein), (D) superoxide dismutase (SOD) activity (U/mg protein), (E) Dot blot analysis of protein carbonyls in liver tissue homogenates and its quantitation shown in (F). (* p < 0.05; ** p < 0.01; **** p < 0.0001; one-way ANOVA followed by Sidak post-hoc multiple comparison test).

### 3.4. Increased methylglyoxal-derived AGE levels in the livers of Glo-1 deleted mice

We then examined the effect of APAP treatment on Glo-1 expression in WT mice by western blot (Figure 5). In WT mice, high dose of APAP reduced the expression of Glo-1 by ∼50% (Figure 5A, 5B). Significant reduction in Glo-2 level was also observed in APAP treated WT and GKO mice (Figure 5B). Treatment with ψ-GSH was able to restore Glo-2 expression to levels in respective saline controls. Compromised Glo pathway in WT after APAP overdose could partly explain equal efficacy of ψ-GSH in WT and GKO mice (Figures 2, 3).

**Figure 5.**
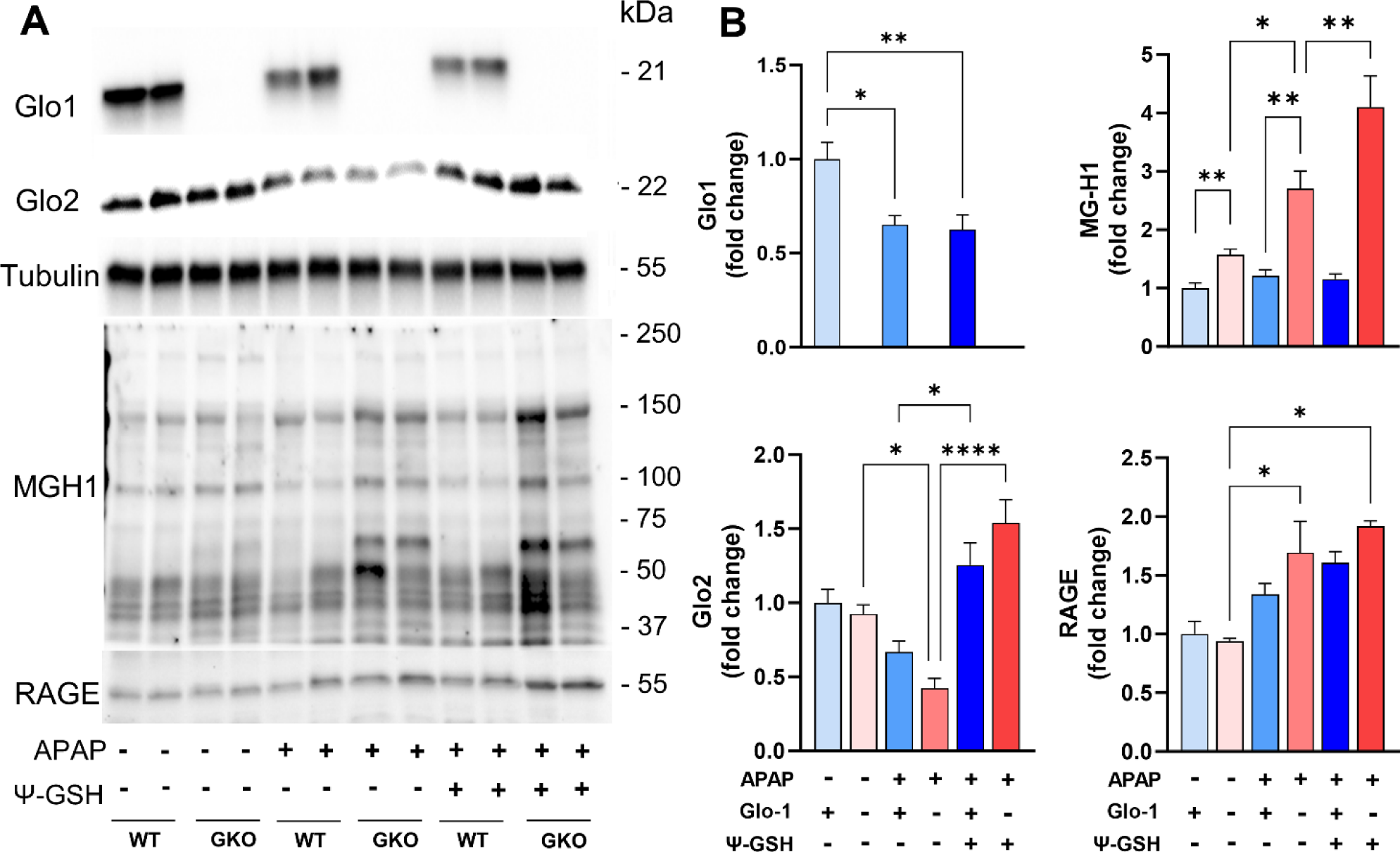
Deletion of Glo-1 increased levels of MEG and MEG-derived AGEs in the livers of mice treated with APAP. (A) Western blot analysis of Glo-1 and Glo-2 expression, MEG-derived AGEs (MG-H1) content and expression of RAGE (receptor of AGE). (B) Quantification of the blots shown in (A). Glo-1 related pathway markers, AGE and RAGE, were significantly elevated in GKO mice livers compared to corresponding WT mice. (* p < 0.05; ** p < 0.01; *** p < 0.001; **** p < 0.0001; one-way ANOVA followed by Tukey’s post-hoc test).

Glo-1 is involved in detoxification of MEG using the cofactor GSH. MEG reacts covalently with lysine and arginine residues in proteins, resulting in formation of irreversible AGEs. It is thus expected that deletion of Glo-1 would increase MEG levels and thus, resultant AGE content in GKO mice. Measurement of AGE levels in the liver tissues of GKO mice has been reported previously by Jang et al.[17]. Western blot analysis using MG-H1 antibody, specific to AGEs derived from MEG, showed significant accumulation of AGEs in APAP-treated GKO livers compared to treated WT liver homogenates (Figure 5A, 5B). Significant difference in the baseline AGE levels of GKO and WT livers was also noted with MEG-specific MG-H1 antibody (Figure 5B), in agreement with the previous report [17]. We further confirmed western blot findings using a commercial mouse AGE ELISA kit (Supporting Information, Figure S5). Baseline liver AGE content in saline treated GKO (14.28 ± 1.87 ng/mg protein) and WT mice (12.47 ± 3.24 ng/mg protein) was similar. AGE content in GKO-APAP and WT-APAP livers (42.56 ± 3.53 *vs* 34.10 ± 4.32 ng/mg protein, respectively) were significantly elevated in comparison with corresponding saline-treated controls. Measurement of AGE levels using ELISA assay corroborated western blot data and showed highest elevation of AGE levels in GKO-APAP, suggesting contribution of AGE in exacerbation of APAP toxicity. Treatment with ψ-GSH was ineffective in reducing AGE content in both WT and GKO mice.

The toxicity induced by AGEs is believed to be through their interaction with receptor for AGEs (RAGE), which triggers various signaling cascades leading to further inflammation and oxidative stress. It also plays significant role in cell death signaling in various pathologies. Thus, we examined the level of RAGE in the livers of APAP treated animals. Western blot analysis showed significantly increased expression of RAGE levels in APAP treated groups (1.5-1.8 fold), with highest levels found in APAP-GKO mice (1.8-fold) compared to saline treated WT controls, as shown in quantitation in Figure 5B. This data suggests that Glo-1 deletion leads to elevation of AGE content upon APAP insult and thus, resultant cellular toxicity through AGE-RAGE interaction. However, ψ-GSH is unable to correct AGE/RAGE after acute treatment, possibly due to compromised Glo-1 expression after APAP overdose. This further supports our notion that Glo-1 modulates hepatotoxicity of APAP, as indicated by the increased expressions of individual components of Glo-1/AGE/RAGE axis.

### 3.5. Distinct cell death mechanisms activated by high-dose APAP in the absence of Glo-1

Given the importance of Glo-1 in detoxification of MEG, inhibition of Glo-1 has been used by us and others for development of anticancer therapeutics [37–40]. Glo-1 inhibition depresses cell proliferation and induces apoptosis in cancer cells, the degree of which is dependent on the concentration of generated MEG. Knockout of Glo-1 in human pluripotent cells (hiPSCs) and liver cell line caused mitochondrial impairment, including diminished membrane potential and dampened respiratory function [41]. In this study, histological analysis (Figure 2) of APAP treated GKO mouse liver displayed increased microvesicular steatosis, most commonly associated with mitochondrial dysfunction. Additionally, ballooning (swollen) hepatocyte degeneration was apparent in the GKO-APAP mice, which is a sign of lipotoxic liver injury. This suggests activation of distinct cell death mechanisms in GKO mice upon high-dose APAP treatment. We therefore investigated the effect of Glo-1 pathway on APAP-induced cell death. We studied the effect of APAP treatment and Glo-1 deletion on the expression of apoptosis (Bax and caspase-3), autophagy (p62) and necrosis markers (HMGB-1, RIPK3) (Figure 6A). Levels of Bax protein were elevated after APAP treatment, with higher levels found in APAP-GKO group (2.1-fold over GKO-saline and 1.75-fold over WT-APAP, p < 0.05 *t-*test; Figure 6B). In order to determine consequences of Bax upregulation, we examined the apoptosome marker caspase-3. Although intact procaspase-3 could not be detected in these samples, cleaved caspase-3 was visible in the WT-APAP group, but only upon high exposure, and not in the APAP-GKO group. This observation coupled with absence of apoptotic morphology in APAP treated groups suggests minimal contribution of apoptosis in APAP-induced cell death, irrespective of Glo-1 expression. This observation supports general knowledge in the field that describes APAP-induced hepatotoxicity does not involve apoptotic cell death [42] and caspase inhibitors are ineffective as therapeutic antidotes. Similarly, analysis of the autophagy substrate p62 showed higher accumulation of p62 protein in APAP-treated samples in both WT and GKO mice (3.4-3.8 fold compared to corresponding saline controls), indicating inhibition of autophagy by high-dose APAP. Experimental NAFLD models and human patients display compromised autophagy and thus, are suggested to have higher-risk of APAP-induced liver toxicity [43].

**Figure 6.**
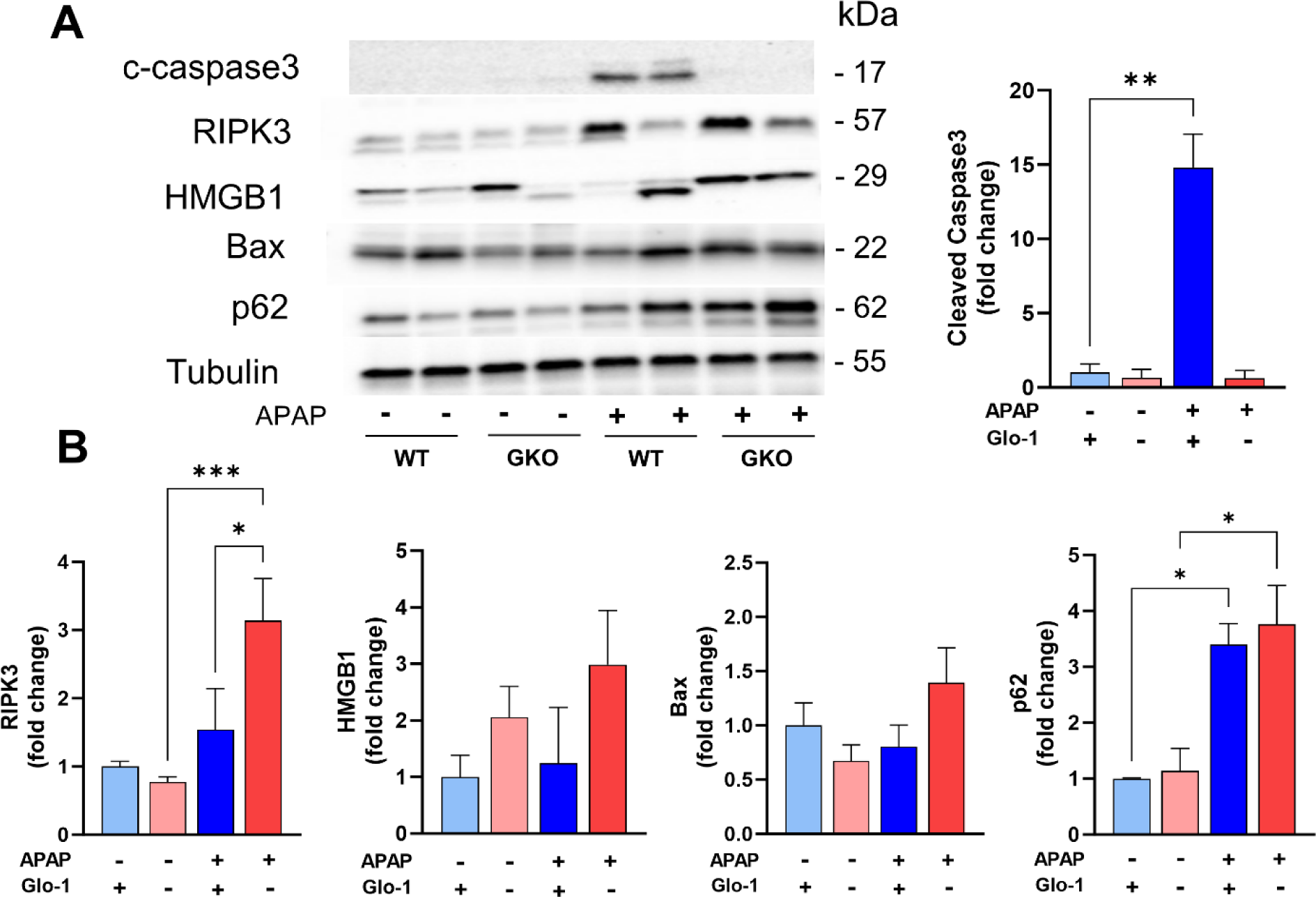
Distinct cell death mechanisms activated by high dose APAP in the absence of Glo-1. (A) Western blot analysis of cleaved caspase3 (c-caspase3), RIPK3, HMGB-1, Bax, and p62. (B) Quantification of blots shown in (A). (* p < 0.05; ** p < 0.01; *** p < 0.001; **** p < 0.0001; one-way ANOVA followed by Tukey’s post-hoc test).

The consensus in the field is that APAP liver toxicity stems largely from necrosis or necroptosis [42]. We examined the expression of necrosis and necroptosis markers: high-mobility group box 1 (HMGB1) and receptor interacting protein kinase 3 (RIPK3), respectively. Different isoforms of HMGB1 were observed in APAP treated livers in the presence and absence of Glo-1, which could indicate different conditions of its release, including different post-translational modifications or proteolytic cleavage. These modifications could contribute differently toward inflammatory response generated by HMGB1. Indeed, loss of integrity of plasma and nuclear membrane results in release of pro-inflammatory form of HMGB1. Higher inflammation seen in WT-APAP livers indicate early release of HMGB1 in these mice [44], which corresponds to higher expression of cleaved HMGB1 compared to WT-saline group. Intensity of HMGB1 bands was higher in saline treated GKO livers, which is further elevated upon exposure to high-dose APAP. Change in HMGB1 isoform released in APAP treated GKO and WT mice, possibly due to proteolytic cleavage, is expected to affect its translocation and release, and could be responsible for differences in the inflammatory response. We further examined RIPK3 expression levels, which is reported to be a molecular switch between apoptosis to necroptosis in APAP hepatotoxicity [45]. APAP-treatment increased RIPK3 protein levels in both WT and GKO mice, while the highest increase was seen in GKO-APAP mice compared to GKO-saline mice (4.1-fold) and WT-APAP group (2.1-fold). HMGB1 is known as a prototypic damage-associated molecular patterns (DAMP) by engagement of cytokines and cell surface receptors such as RAGE. It is implicated in amplification of hepatocyte necrosis via RIPK3 dependent pathway [46]. It is also interesting to note that accumulated p62 has been shown to interact with RIPK1 and RIPK3, facilitating the creation of necrosomes, and ultimately, necroptosis [47].

## 4. Discussion

In this study, we examined the role of Glo-1 in acute APAP overdose-induced liver injury using a constitutive Glo-1 deleted mouse model. We also examined the effect of a synthetic Glo-1 cofactor surrogate and general antioxidant, ψ-GSH on this model. Our results indicate that mice lacking Glo-1 expression show distinct pattern of hepatotoxic effect after APAP overdose when compared to the WT mice. This finding is corroborated by elevated oxidative stress, APAP metabolism, liver chemistry/histopathology and activation of distinct cell death pathways. This change caused by APAP overdose was more prominent in male compared to female mice. Lower susceptibility of female mice toward APAP injury has been reported previously, with the potential cause being improved detoxification or repair mechanisms and potential role of estradiol [48]. Our study displayed higher levels of the non-toxic APAP-sulfate, APAP-GSH and APAP itself in female WT and GKO mice. This would indicate effective conjugation and clearance of intact APAP in females compared to males being at least partly responsible for their resistance to APAP hepatotoxicity.

Glo-1, a cytosolic enzyme, is a part of the glyoxalase system, responsible for the detoxification of MEG, which provides protection against dicarbonyl stress [49]. Stress induced by APAP overdose further augmented the effects of Glo-1 deletion and significantly aggravated the levels of AGEs when compared to corresponding WT mice. High concentrations of AGEs are known to activate RAGE, which is promoted by oxidative stress and inflammation and have consequences in various liver disorders [50]. Indeed, expression of RAGE was more pronounced in GKO-APAP group compared to respective saline treated controls. Interaction of AGE-RAGE is expected to contribute toward propagation and acceleration of APAP-induced toxic effects. Consequently, hallmark oxidative stress markers such as lipid peroxidation, protein carbonyl and oxidized GSH levels were elevated in GKO-APAP mice. High oxidative stress in GKO mice limited efficient detoxification of APAP in GKO male mice, as seen by reduced levels of non-toxic APAP adducts with GSH and Cys, as well as sulfated APAP metabolites. Treatment with the Glo-1 cofactor, ψ-GSH, successfully mitigated ALT elevation and biochemical consequences of APAP overdose, irrespective of Glo-1 status. Compromised Glo-1 expression in APAP treated WT mice, compensatory mechanisms in GKO mice and the Glo-1 independent antioxidant capability of ψ-GSH could account for the lack of Glo-1 dependence for ψ-GSH’s pharmacological action. Histopathological examination revealed striking changes in liver lipid accumulation in GKO livers, with hepatocyte degeneration caused by microvesicular steatosis. This observation suggests distinct degenerative pathology induced by Glo-1 deletion, which further intensifies APAP-induced liver toxicity. Increased AGE content with RAGE stimulation have also been found in liver steatosis and fibrosis in the SD rats [50] and in NAFLD patients. It also induces inflammation, proliferation and fibrosis in the hepatic stellate cells (HSCs) by stimulating TGF-β1 expression [51]. Absence of functional Glo-1 also elevated necrosis, specifically necroptosis-induced cell death and diminished signs of apoptosis as evident by increased HMGB1 and RIPK3 levels and absence of caspase-3 processing in APAP treated GKO mice. Time-course of APAP toxicity in these mice could help delineate contribution of Glo-1 in initiating various cell-death cascades. Differences in WT and GKO mice support contribution of Glo-1 toward necrosis since its deletion shows activation of programmed necrosis (necroptosis). Liver steatosis in GKO-APAP group could trigger lethal lipotoxic signals, initiating hepatocyte degeneration, mitochondrial dysfunction, and fibrotic reactions leading to NAFLD-type pathology [52], enhancing hepatotoxic effects of APAP.

Effective address to deleterious APAP effects is provided by the clinical antidote, NAC, by correcting the tissue redox status and quenching the reactive metabolite of APAP. However, its use is hampered by a narrow treatment window, gastrointestinal side effects and anaphylaxis. Other means to alleviate oxidative stress, thus could lend therapeutic approaches to mitigate the hepatotoxicity of APAP poisoning. Exogenous administration and overexpression of antioxidant enzymes, SOD and glutathione peroxidase, have provided promise in mitigating APAP toxicity in preclinical studies [53,54]. Blockage of RAGE also leads to attenuation of APAP-induced liver damage [6]. Diabetes/hyperglycemia known to activate AGE/RAGE signaling is considered one of the risk factors for acute liver injury [55]. Thus, regulating the consequences of toxic glucose metabolites such as AGE formation by balancing cellular redox potential and inhibition of RAGE, rather than Glo-1 function restoration, would be expected to lend an effective therapeutic strategy. Supplementation with bioavailable GSH analogs or precursors have shown promise in preclinical and clinical setting [13,56]. Apart from their innate antioxidant potential, such compounds are expected to engage GSH-dependent enzymes influential in restoring redox homeostasis. The results of this study highlight importance of one such GSH enzyme, Glo-1, in modulating hepatotoxic effects of APAP. Furthermore, characterization of cell death mechanisms underscores regulation of RIPK3-dependent mechanisms for alleviation of APAP-induced liver injury. Additional studies are however needed to understand time- and dose-dependent effects of APAP overdose on Glo-1 pathway intermediates and cell death mechanisms.

## 5. Conclusions

Taken together, our data demonstrates the important function conducted by Glo-1 in liver homeostasis and in modulating glycation-induced oxidative stress resulting from acute toxic insults. Supplementation with an antioxidant GSH mimetic alleviated lethal lipotoxic phenotype displayed by Glo-1 deleted hepatocytes and also prevented oxidative stress and hepatocyte degeneration. Such intervention could have application in treatment of acute liver toxicity such as that induced by extraneous agents or chronic liver illness leading to NAFLD or NASH pathology. Efforts to understand time course of liver toxicity progression after APAP overdose, and factors responsible for resistance of female mice toward APAP toxicity are currently underway. This study adds greatly to our understanding of the Glo-1/AGE/RAGE axis in APAP-induced hepatotoxicity and emphasizes the importance of mitigating redox imbalance to effectively address the initial stages of acute liver injury.

## Supplementary Materials

The following supporting information can be downloaded at: XXXX, Supplementary File 1 (Supplementary Figures S1-S5 and Table 1).

## Author Contributions

Conceptualization, S.S.M. and M.K.L.; performing experiments, P.D., S.R., W.X., J.X.; data collection and analysis, P.D., S.R., W.X., M.K.L., S.S.M.; writing—original draft preparation, P.D., S.S.M.; writing—review and editing, M.K.L, R.V., D.M.S., S.R., W.X., J.X.; supervision, S.S.M. and M.K.L.; funding acquisition, S.S.M., R.V., and M.K.L. All authors have read and agreed to the published version of the manuscript.

## Institutional Review Board Statement

The study was conducted in accordance with the Declaration of Helsinki, and approved by the Institutional Animal Care and Use Committee (IACUC) at the University of Minnesota (protocol ID 2205-40054A and date of approval 07/12/2022). All applicable ethical standards required by University of Minnesota IACUC were followed.

## Data Availability Statement

Chemical compounds and all raw data or original images will be made available upon request made to More.

## Acknowledgments

The authors thank Dr. Emma Torii, Comparative Pathology Shared Resource Masonic Cancer Center, University of Minnesota, for processing of the liver tissues for histopathological analysis and for valuable discussions. We also thank Dr. Indrani Poddar and Ms. Rachel Tappe for maintaining and generating the GKO mouse cohorts. Graphical abstract was created with BioRender.com.

