## Supplementary File 1 for "Deletion of Glyoxalase 1 exacerbates acetaminophen-induced hepatotoxicity in mice"

Supplementary Figures S1-S5 and Table 1

For

**Deletion of Glyoxalase 1 exacerbates acetaminophen-induced hepatotoxicity in mice**

Prakashkumar Dobariya^1,†^, Wei Xie^1,†^, Swetha Pavani Rao^1^, Jiashu Xie^1^, Davis M. Seelig^2,3^, Robert Vince^1^, Michael K. Lee^4,5^, and Swati S. More^1,*^

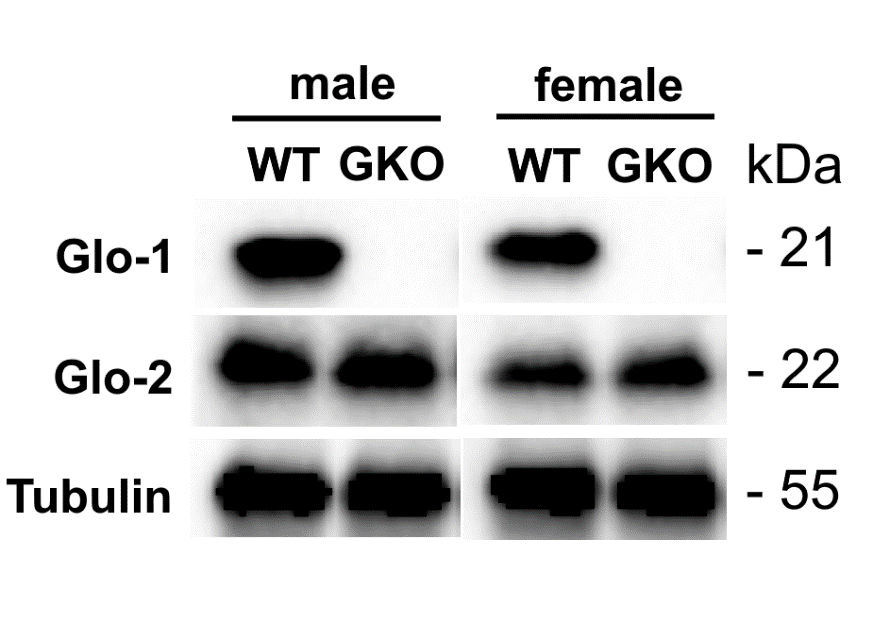
**Figure S1.** Characterization of GKO mice by western blot for expression of Glo-1 and Glo-2. Deletion of Glo-1 was confirmed in GKO mice by the absence of protein band.

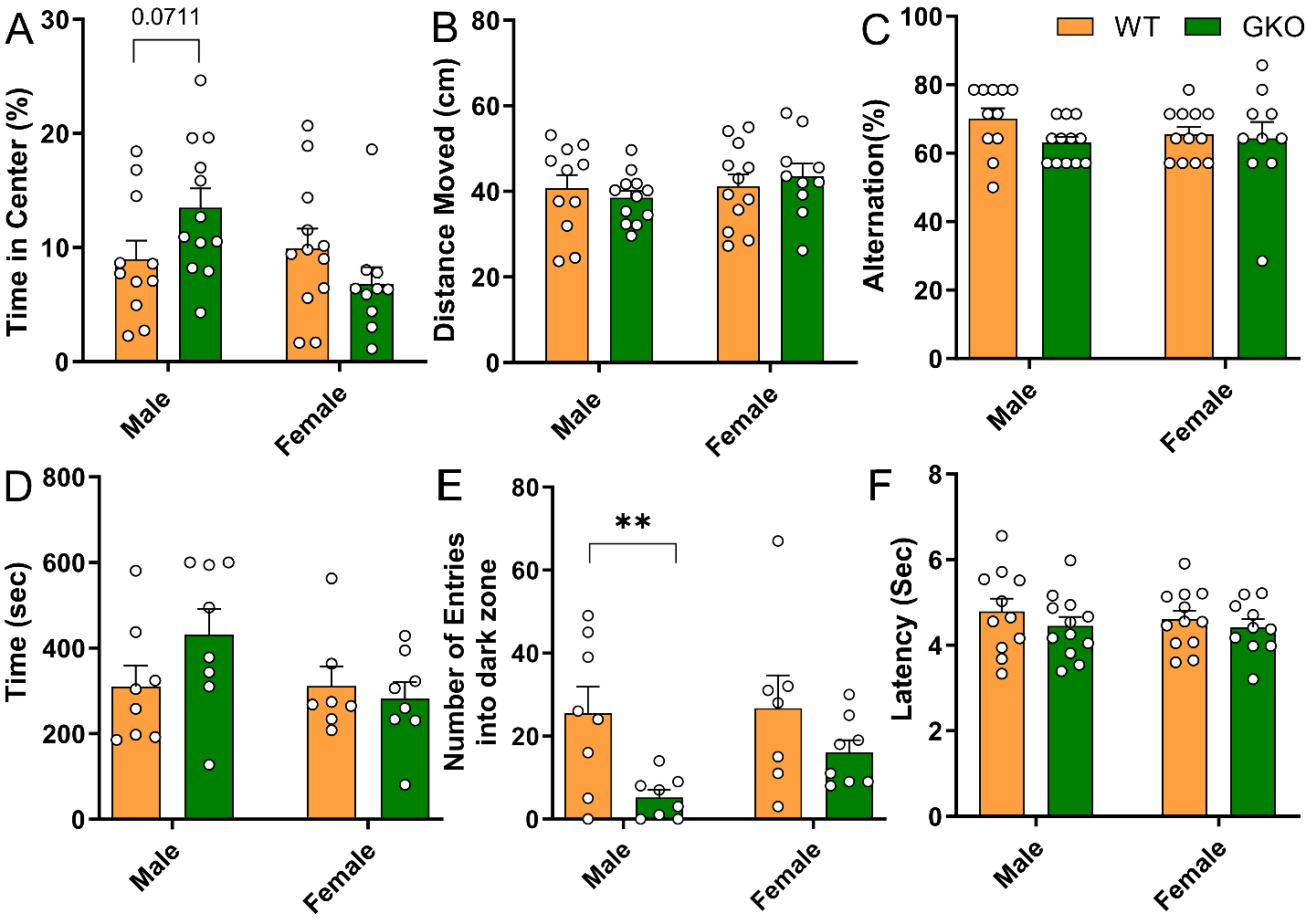

**Figure S2.** Deletion of Glo-1 reduces anxiety-like behavior. (A-B) Open field test showing time spent in the center zone (A) by age-matched WT ( ) and GKO mice ( ); (B) total distance moved during the open filed test; (C) T-maze spontaneous alternation test showing %alternation rate; (D-E) Light-dark box test showing time spent in light zone (D) and number of transitions (E) during the testing period. (F) Tail flick test showing the latency of tail flick response. (n = 11-12 per group). Data are shown as mean ± SEM (* p < 0.05 by two-way ANOVA using Sidak multiple comparison post-hoc test).

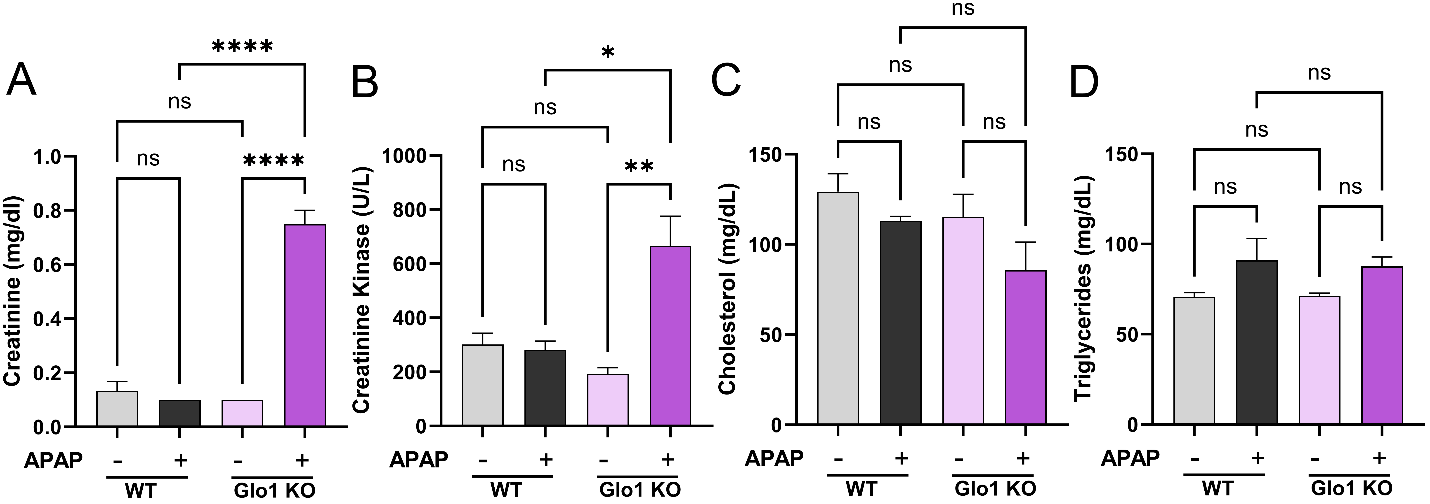

**Figure S3.** Analysis of serum creatinine (A), creatinine kinase (B), cholesterol (C) and triglyceride (D) levels of WT and GKO mice treated with high-dose APAP. Data are shown as mean ± SEM (*p < 0.05, ** p<0.01, **** p<0.0001 by one-way ANOVA using Sidak multiple comparison post-hoc test).

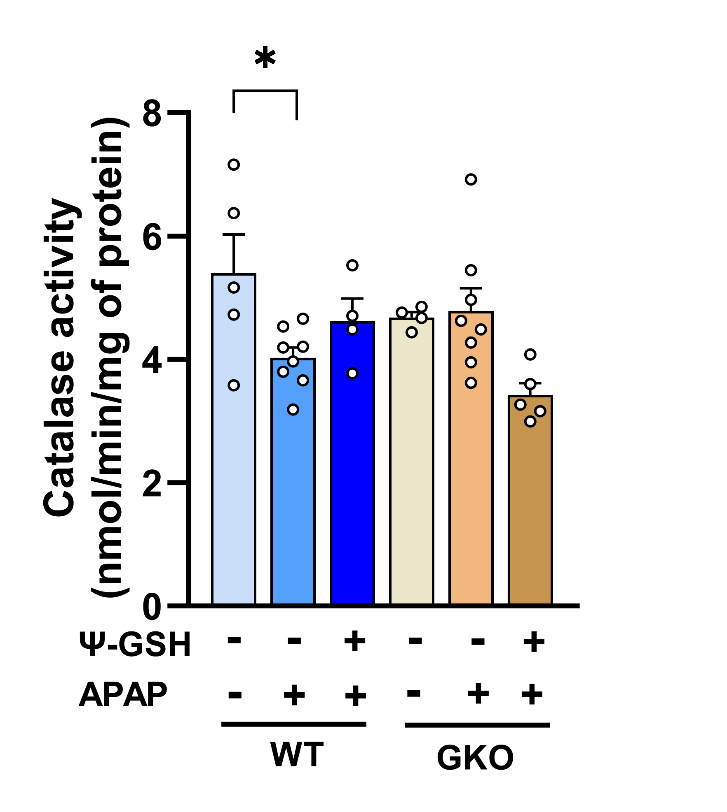

**Figure S4.** Liver catalase activity of mice treated with APAP. Except for the reduced catalase activity in WT-APAP group compared to WT-saline, no significant differences were noted within different treatment groups. Data are shown as mean ± SEM (* p < 0.05, one-way ANOVA followed by Tukey’s multiple comparison post-hoc test).

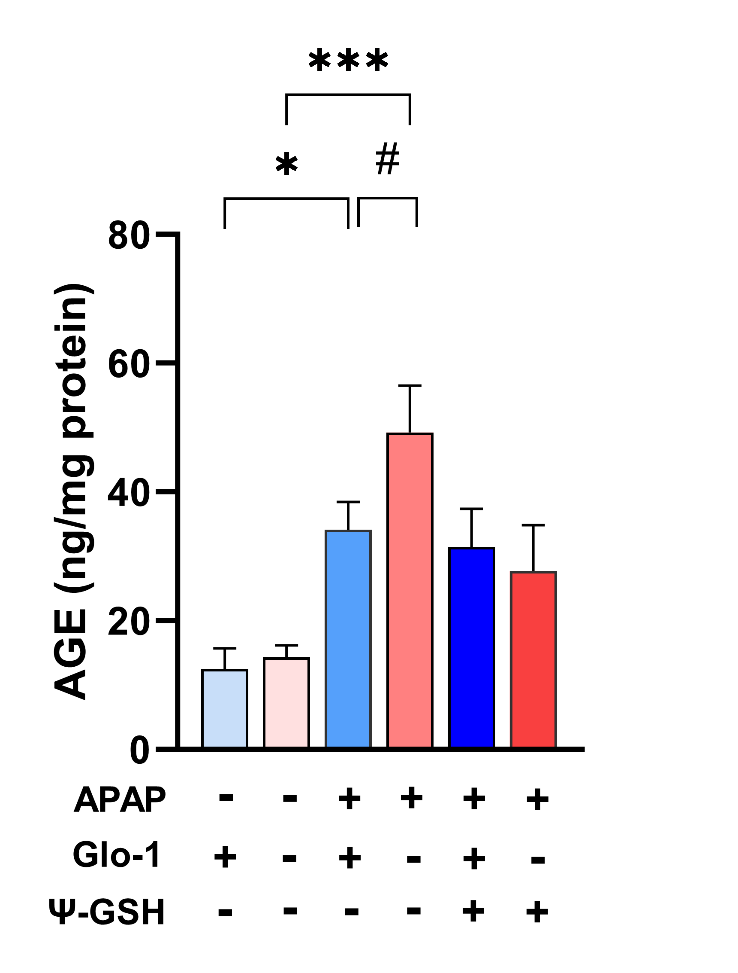
**Figure S5.** Quantitation of liver AGE content by a commercial mouse AGE-ELISA. APAP treatment increased AGE content in WT and GKO livers. Ψ-GSH treatment was unable to reduce AGE levels. Data are shown as mean ± SEM (* p < 0.05, *** p < 0.001, one-way ANOVA followed by Tukey’s multiple comparison post-hoc test; # p = 0.05, *t*-test)

**Table 1.** Concentration of APAP metabolites in serum samples expressed as nmol per μL serum

| **Metabolite** | **Wild Type** | **GKO** | **Wild Type** | **GKO** |
| --- | --- | --- | --- | --- |
|  | **Male** | | **Female** | |
| APAP | 690.88 ± 147.57 | 409.45 ± 132.56 | 1476.31 ± 538.24 | 2370.11 ± 565.77 (0.0150* *vs* GKO male) |
| APAP-glucuronide | 1415.64 ± 678.44 | 671.09 ± 161.73 | 3019.23 ± 1586.09 | 7347 ± 1880.66 (0.0459* *vs* WT male; 0.0168 *vs* GKO male) |
| APAP-sulfate | 230.12 ± 41.26 | 65.99 ± 36.01  (0.0241* *vs* WT male) | 1987.66 ± 338.46  (0.0021** *vs* WT male) | 2343.81 ± 330.71  (0.0005*** *vs* GKO male) |
| APAP-GSH | 28.08 ± 8.25 | 6.147 ± 3.03 (0.0187* *vs* WT male) | 34.48 ± 22.47 | 41.15 ± 4.17  (0.0001*** *vs* GKO male) |
| APAP-Cys | 916.57 ± 338.22 | 150.19 ± 32.66 (0.065 *vs* WT male) | 561.63 ± 251.19 | 352.31 ± 118.89 |
| APAP-NAC | 427.30 ± 73.82 | 88.28 ± 24.67 (0.0048** *vs* WT male) | 637.24 ± 347.69 | 250.43 ± 79.79 |
| APAP-GSH | 28.08 ± 8.25 | 6.147 ± 3.03 (0.0187* *vs* WT male) | 34.48 ± 22.47 | 41.15 ± 4.17  (0.0001*** *vs* GKO male) |
| APAP-OMe | 1.96 ± 1.08 | 0.19 ± 0.22 (0.0184* *vs* WT male) | 2.66 ± 1.11 | 4.01 ± 1.58 (0.0030** *vs* GKO male) |
